# Postmating sexual selection and the enigmatic jawed genitalia of *Callosobruchus subinnotatus*

**DOI:** 10.1101/116731

**Authors:** Merel M. van Haren, Johanna Liljestrand Rönn, Menno Schilthuizen, Göran Arnqvist

## Abstract

Insect genitalia exhibit rapid divergent evolution. Truly extraordinary structures have evolved in some groups, presumably as a result of post-mating sexual selection. To increase our understanding of this phenomenon, we studied the function of one such structure. The male genitalia of *Callosobruchus subinnotatus* (Coleoptera: Bruchinae) contain a pair of jaw-like structures with unknown function. Here, we used phenotypic engineering to ablate the teeth on these jaws. We then experimentally assessed the effects of ablation of the genital jaws on mating duration, ejaculate weight, male fertilization success and female fecundity, using a double-mating experimental design. We predicted that copulatory wounding in females should be positively related to male fertilization success. However, we found no significant correlation between genital tract scarring in females and male fertilization success. Male fertilization success was, however, positively related to the amount of ejaculate transferred by males and negatively related to female ejaculate dumping. Ablation of male genital jaws did not affect male relative fertilization success but resulted in a reduction in female egg production. Our results suggest that postmating sexual selection in males indeed favors these genital jaws, but not primarily through an elevated relative success in sperm competition but by increasing female egg production.

## Introduction

Insect genitalia exhibit rapid divergent evolution (Hosken and Stockley, 2004; Eberhard, 2004; Eberhard, 2010). There is now little doubt that this is due to postmating sexual selection (Birkhead and Pizzari, 2002; Hosken and Stockley, 2004; Arnqvist, 2014), generated either by conventional cryptic female choice (CFC) whereby female traits are evolving to gain benefits (Eberhard, 2006) or by sexually antagonistic coevolution (SAC) whereby female traits are evolving to minimize direct costs imposed by males (Arnqvist and Rowe, 2005). This revolutionary process can result in the evolution of remarkable structures, such as prominent sclerotized structures of male genitalia that causes injuries to females. The function of these structures have only rarely been addressed, but can involve enabling copulations (Grieshop and Polak, 2012) or increasing male fertilization success by allowing passage of male seminal fluid (Kamimura, 2010; Hotzy et al., 2012).

Seed beetles are widely employed in studies of postcopulatory sexual selection and are well known for showing harmful male genital structures (Hotzy et al., 2012; Rönn et al., 2007; Sakurai et al., 2012) that damage the female copulatory tract. *Callosobruchus subinnotatus* (Coleoptera, Bruchinae) is a seed beetle with a particularly interesting male genital morphology, as males are equipped with a pair of prominent sclerotized “jaws” (Fig. 1).

**Fig. 1.**
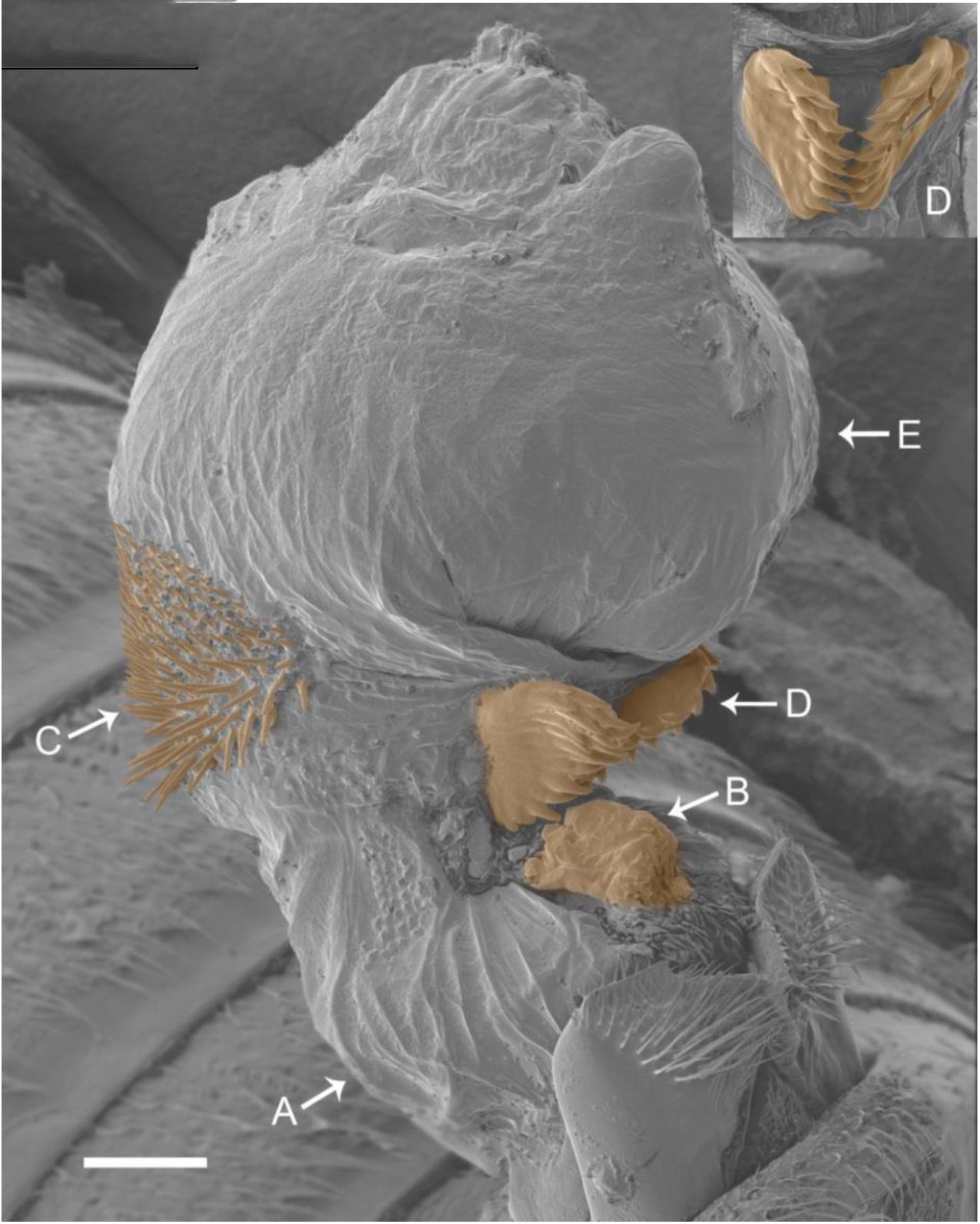
The remarkable male genitalia of *C. subinnotatus.* During copulation, the genitalia unfolds which results in a reformation of its armature. This starts with the expansion of the base (A) of the endophallus. On top of this base sits a sclerotized structure, the basal structure (B), that appears as a thickened fold of the base of the endophallus. At this point, the dorsal spines (C) are clearly visible. The jaw-like structures on the ventral side (D) then join up, due to the expansion of the internal sac tip (E). At this point the jaws are closed and their position appear fixed. The endophallus is distinct from that in other seed beetle species (Rönn et al. 2007). The figure shows an endophallus fixated by critical point drying to prevent tissue from collapsing. Scale bar represents 100μm.

To better understand the evolution of such genital structures, we performed a series of experiments aimed at unveiling the ultimate function of these genital jaws. The jaws clearly cause injury to females: the copulatory duct is abraded or even pierced by the jaws, leaving a v-shaped pattern of melanized scars (Fig. 2).

**Fig. 2.**
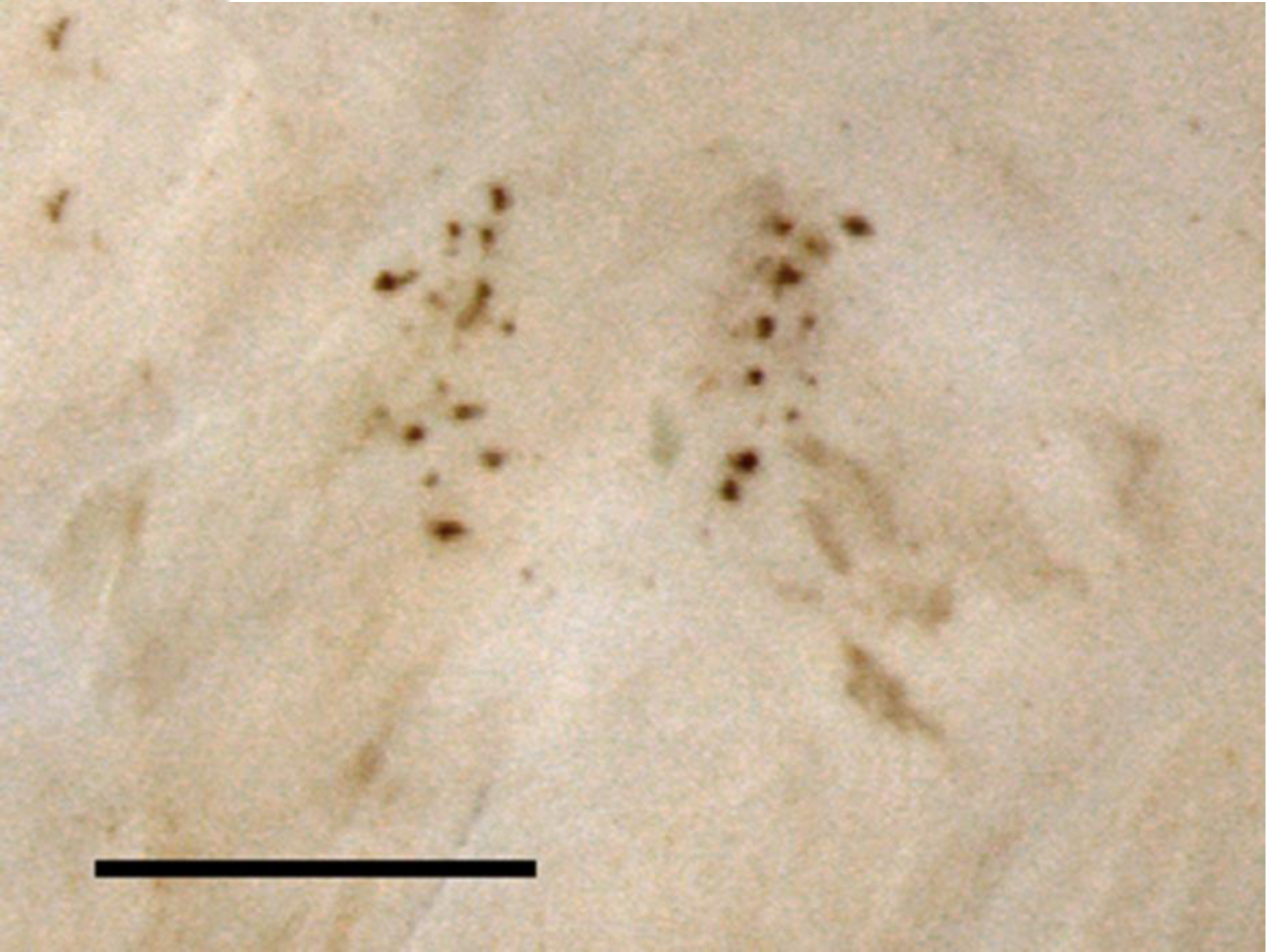
The V-shaped scarring pattern caused by the jaw-like structures in the copulatory duct of females. Scale bar is 200μm.

We hypothesized that the genital jaws may either (1) serve as a holdfast device or (2) may elevate male fertilization success by other means, as it is the case in the closely related species *C. maculatus* (Hotzy and Arnqvist, 2009; Hotzy et al., 2012). Here, we used phenotypic engineering to experimentally manipulate this structure. The paired genital jaws bear spiny teeth-like protrusions (Fig. 1) which formed the target of our manipulation: to smoothen the teeth by abrasion. A complete removal of the jaws would have been interesting, but was impossible as it would have caused detrimental hemorrhage.

## Materials and Methods

Beetles were mass cultured in the laboratory on a 12:12 h L:D photoperiod, 55% RH and a temperature of 29°C in 1000 mL glass bottles (N=3), containing 250ml black eyed-beans (*Vigna unguiculata*) per generation. New generations were set by mixing beetles from each of the jars, to avoid inbreeding (Appleby and Credland, 2001). To generate virgin individuals, beans with eggs were isolated individually in 24 well tissue culture plates. Beetles used in the experiments described below were all of < 48 hours adult age and were kept individually under aphagy in aerated 5ml Eppendorf tubes prior to the experiment.

### Treatment of males

To assess the function of the jaw-like structures, their teeth were smoothened manually following the eversion of their genitalia (Fig. 3).

**Fig. 3.**
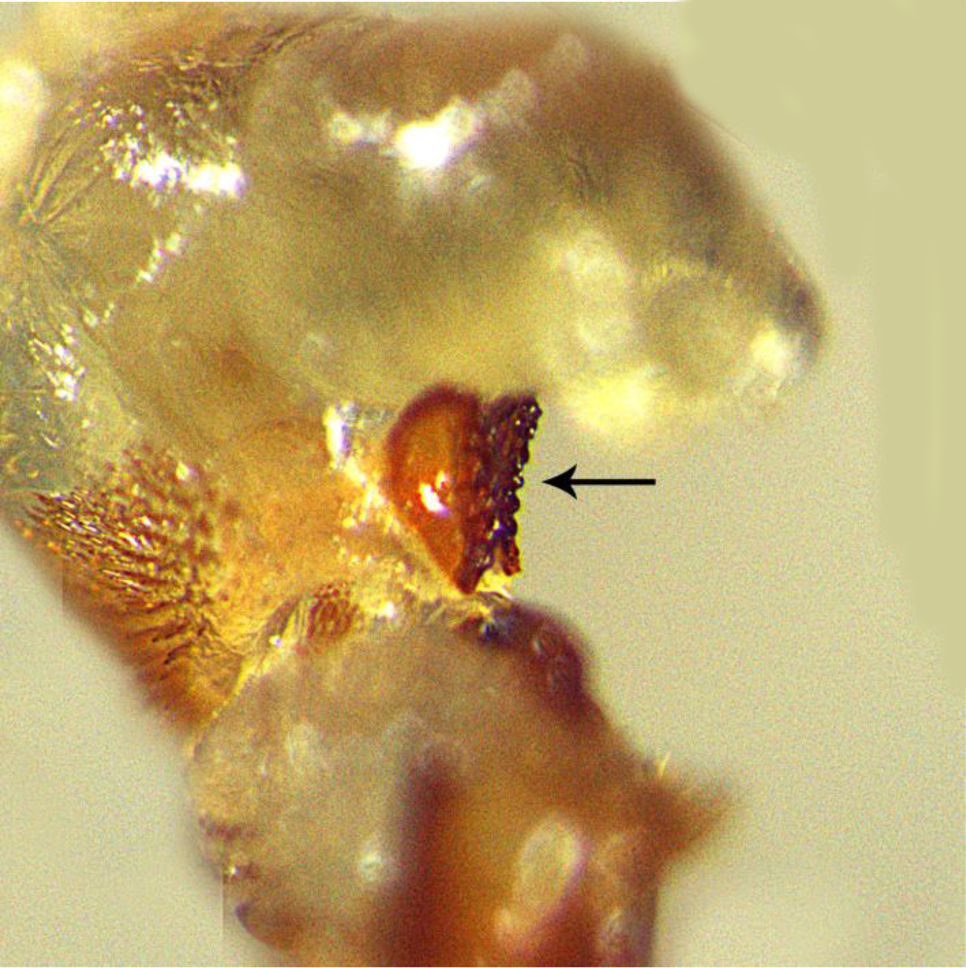
The endophallus of a *C. subinnotatus* male with smoothened teeth. Scale bar is not available, but for references the jaws are approximately 100μm.

The treatment was performed with a file made of a dentist drill (Two striper L201MF3) attached to a probe. To smoothen the teeth, the jaws of lightly anesthetized (CO_2_) males were held in position with a forceps (SS 11200-33 Dumoxel®-Biology CE) (Fig. 4) under a dissecting microscope. All male treatments (see below) were performed with the same method and materials.

**Fig. 4.**
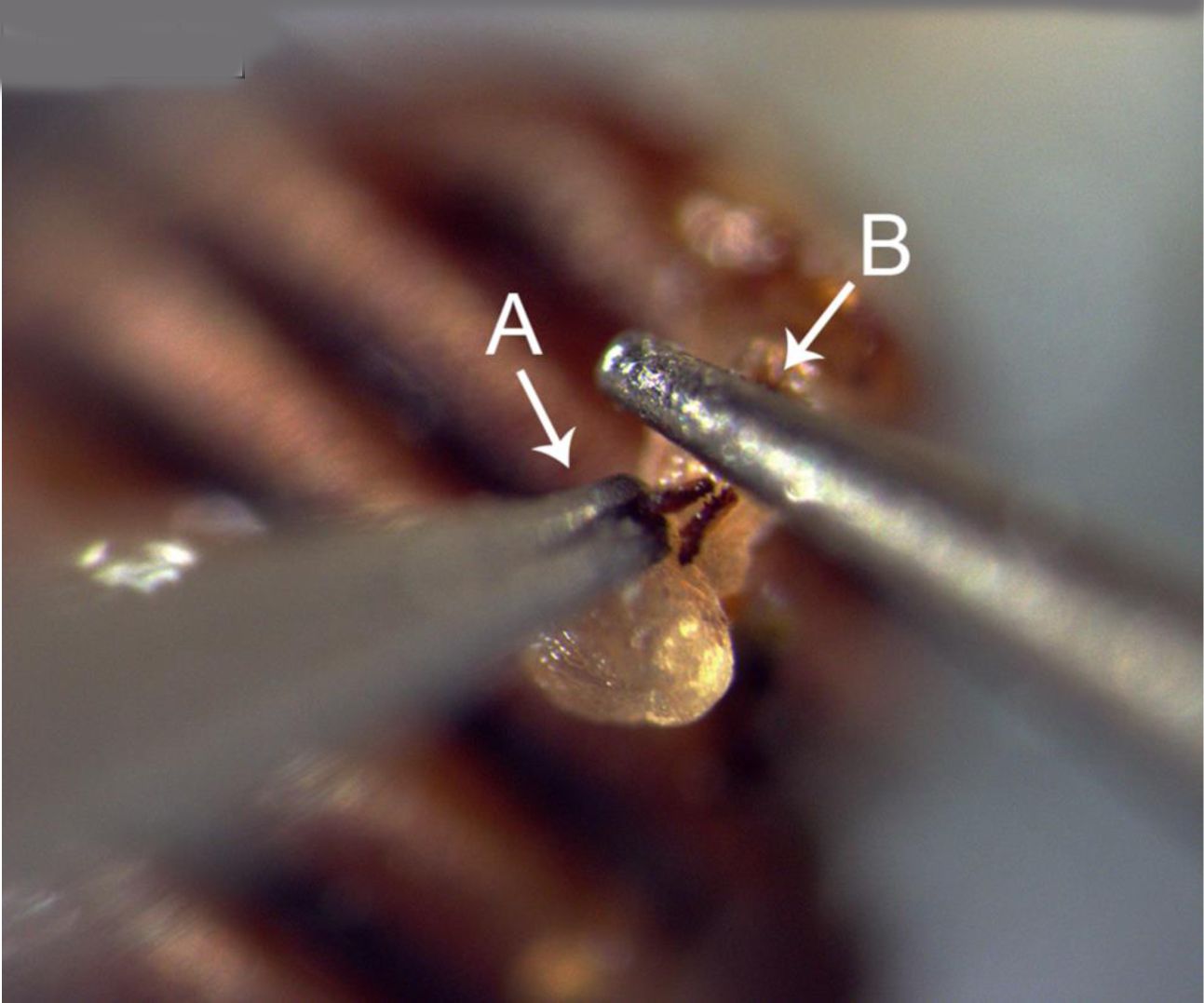
The manipulation of the jaws. A male with everted endophallus is fixed with its elytra on blue-tack. The jaws are held in position with a forceps (A) and then filed down using a dentist drill (B). Scale bar is not available, but for references the jaws are approximately 100μm.

Our experimental design included four treatment groups (see Figure 5 for sample sizes). [A] Some males had the teeth of their genital jaws smoothened - we refer to this as the ablated jaws males (AJ). Three different control groups were created. [B] One group of males were not manipulated in any way, but were left untouched (Non). This group controls for potential effects of CO_2_ anesthesia and genital eversion. To control for ablation per se, two additional control groups were created. [C] One group were treated as AJ males in every respect but instead had another structure of their genitalia ablated, namely the right paramere (APa). [D] The final group of males were also treated as AJ males in every respect but served as a surgical control in the sense that they had a non-genitalic structure ablated, namely the rim of the pygidium (APy) (Fig. S1).

**Fig. 5.**
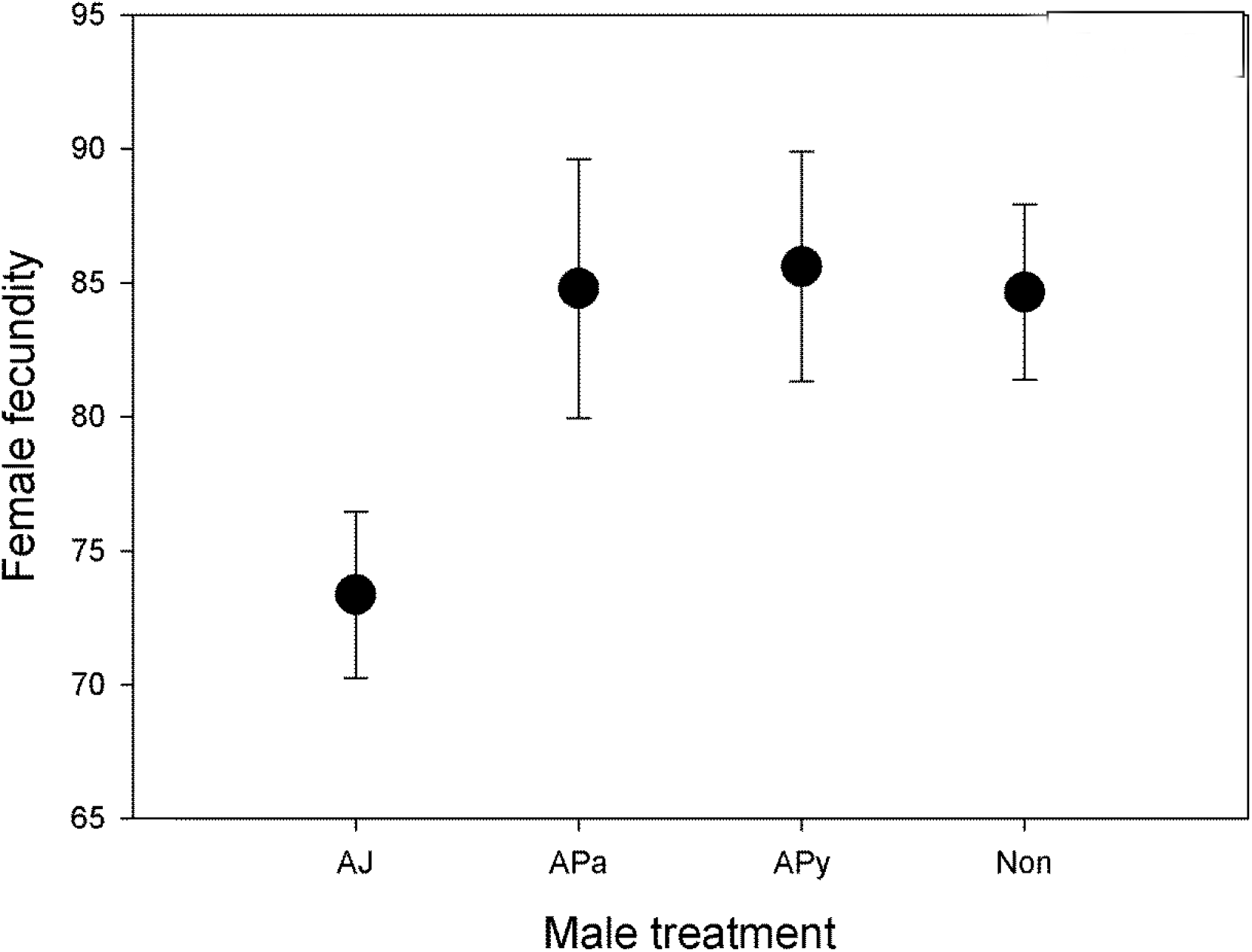
The total number of eggs laid by females. Females laid fewer eggs if one of her mates had ablated genital jaws (AJ) (GLM: *F*_3,148_ = 3.01, *P* = 0.032. *N=55* for AJ, *N*=23 for APa, N=29 for APy, and *N*=50 for Non), compared to mates from the control groups. Shown is marginal mean (±SE) number of eggs.

Focal males were thus treated 18 hours before they were used in the experiments described below. During the treatment, beetles were lightly anesthetized with CO_2_ up to a period of seven minutes by placing them on a FlyStuff Flypad. Virgin reference males were sterilized by irradiating them with a 100 Gy dose from a cesium-137 source. This sterilization technique has been shown to cause lasting sterility in male seed beetles while not compromising male copulation ability and sperm competitive ability (Eady, 1991; Maklakov and Arnqvist, 2009). After the treatment, males were placed in a 6cm petri dish with access to 5ml sugar water solution to recover.

### Treatment of females

Females resist mating, prior to and during copulations, by kicking males with their hind legs and this might affect male fertilization success (Maklakov and Arnqvist, 2009). To assess the influence of female hind leg kicking on the effects of the above male treatment, we also manipulated all females involved in matings with AJ and Non males (not those mated to APa and APy males). One hour before the mating experiment, half of the females were put on ice where the hind tibia were ablated with micro scissors halfway. This renders females unable to reach, and thus resist, males (Crudgington and Siva-Jothy, 2000; Edvardsson and Tregenza, 2005; Maklakov and Arnqvist, 2009). To control for the effect of ablation, the other half of all females were left having intact hind legs during mating but instead had their hind legs ablated one hour after mating with the focal male.

### Mating experiments

We measured eight different aspects of reproductive response to the genital jaw manipulation: mating duration, male ejaculate weight, female ejaculate dumping, the amount of scarring caused in the female copulatory duct, male offensive sperm competition success, male defensive sperm competition success and female fecundity. Moreover, the effect of female kicking during mating on these responses was assessed.

The above reproductive responses were based on a series of standard double-mating experiments in which both defense (P1) and offense (P2) components of sperm competition success were measured using a standard sterile male technique (Boorman and Parker, 1976; Simmons, 2001). Here, females were mated with two males in succession, one of which was irradiated such that his sperm remain motile and fully able to fertilize eggs but carry lethal mutations that render the eggs inviable and the other male was focal and fertile. Here, P1 and P2 denote the proportion of offspring that is fertilized by the focal male when he is first or second to mate, respectively, with a given female in such a double mating experiment. Briefly, focal experimental males, sterile reference males and females were first weighed on a balance (Sartorius ME235S Genius). Mating couples were then immediately introduced in pairs in 6cm petri dishes and placed in dark climate chambers under rearing conditions, during very early morning which represents the peak mating time *C. subinnotatus* (MBata et al., 1997). The initiation and termination of mating were recorded. Pairs that did not mate within 90 minutes were discarded. After mating, both male and female were weighed a second time. Females were placed individually in 10cm petri dishes with +/−40 beans and access to 5ml sugar water solution and were stored in climate chambers for 48 hours. Following this intermating interval, females were remated to a second male following the same protocol as for the first mating. In the sperm offense assays (P2), the first male was a sterile reference male and the second male was a focal experimental male. In the sperm defense assays (P1), this order was reversed. The petri dish with beans and eggs from the intermating interval was incubated for 10 days in a climate chamber, after which all hatched and unhatched eggs were counted.

Male weight loss during mating provides a measure of male ejaculate weight in these insects (Savalli and Fox, 1998; Rönn et al., 2008). The reduction in male weight during copulation was significantly correlated with the increase in female weight across all matings (*r* = 0.39, *P* < 0.001, N = 326). The fact that the correlation was not stronger is primarily due to partial ejaculate dumping immediately after copulation by females, a phenomenon common in seed beetles (Booksmythe et al., 2014) as well as in insects in general (Perry and Rowe, 2008). In our experiments, mean male weight loss was on average 18.2 × 10^-5^ g and mean female weight gain was on average 12.7 × 10^−5^ g (paired t-test: t_325_ = 8.55, *P* < 0.001), suggesting that females dump some 30% of the ejaculate on average. Here, we thus used male weight loss as a measure of ejaculate weight and the difference between male weight loss and female weight gain as a measure of female ejaculate dumping.

Following the second mating, the females were placed in new petri dishes provided with ca. 40 black-eyed beans and a 5ml Eppendorf tube containing sugar water and was allowed 7 days to lay eggs and heal copulatory injuries. After this time, females were frozen (−21°C). After incubation for another ten days, the petri dishes containing eggs on beans were also frozen, to prevent beetles from hatching. All eggs were subsequently counted and we recorded whether each egg was hatched or unhatched. Female were subsequently thawed and the copulatory duct and the bursa copulatrix was separated from the female abdomen, cut open and placed on a microscopic slide, enclosed in glycerin, and covered with a cover slip. The dissections were performed under a Leica M165C microscope. A photo was taken of the dissected bursa with a motorized Zeiss V20 with MRc5 camera and Axiovision software. The images were subsequently analyzed in ImageJ. The image was adjusted into an 8bit format and a threshold was set to distinguish scar tissue from non-scar tissue. We quantified scarring in females as both the number of scars and the area covered by scars, expressed in pixels. All scars were included, since it is not possible to unambiguously distinguish between scars caused by the genital jaws and other types of genital spines (Fig. 1).

### Statistical analyses

In our main models, we modelled the fertilization/reproductive success of the focal male, using his mating order (first [P1] or second [P2]) as factor. For fertilization success, we employed generalized linear models of the number of hatched eggs, using binomial errors with a complementary log-log link function and an empirically derived dispersion parameter where the total number of eggs laid after the second mating was used as the binomial denominator. Conventional general linear models were used for other inferences. Inferential models included our factorial variables (P1/P2, male treatment, female treatment) and any covariates with noticeable effects. Interactions were only included when statistically significant. Potential covariates included body weight of males and females, ejaculate size, sperm dumping, mating duration, scarring in females and the number of eggs laid by females between matings. Models of the effects of female leg treatment were restricted to include only AJ and Non males (see above). Four females that laid <4 eggs after the second mating were excluded from our data set. In addition, two observations with standardized residuals >4 were excluded from the analyses of scarring in females. Analyses were performed with Genstat v.18 and Systat v.13.

## Results

The overall fertilization success of the last male to mate, i.e. P2, was approximately 0.68 in *C. subinnotatus*. The model predictions, adjusted for covariates, were P1 = 0.39 (SE = 0.03) and P2 = 0.76 (SE = 0.03) but both of these values are likely somewhat inflated as a result of a slight competitive advantage of normal sperm over irradiated sperm.

Our inferential model of variation in male fertilization success (Table 1) was highly significant overall (*F*_11,146_ = 7.73, *P* < 0.001). However, male genital treatment had no significant effect on male fertilization success under sperm competition, measured as the proportion of a female’s offspring fertilized by the focal male, but both ejaculate weights and ejaculate dumping by females was associated with male fertilization success (Table 1). Interestingly, focal male fertilization success increased with his ejaculate weight (*β′* = 0.04, SE_*β*_ = 0.01) and decreased with female ejaculate dumping (*β′* = −0.02, SE_*β*_ = 0.01). We found no significant effect of female leg treatment or any other covariates on male fertilization success.

**Table 1.** Analysis of deviance of a generalized linear model of variation in male fertilization success under sperm competition in our double-mating experiment.

| Source | DF | Deviance | Deviance<br>ratio | *P* (from *F*) |
| --- | --- | --- | --- | --- |
| P1/P2 | 1 | 845.83 | 56.08 | <0.001 |
| Male treatment | 3 | 99.65 | 2.20 | 0.090 |
| P1/P2 × male treatment | 3 | 59.97 | 1.33 | 0.268 |
| Eggs laid between matings | 1 | 26.91 | 1.78 | 0.184 |
| Focal male ejaculate weight | 1 | 103.48 | 6.86 | 0.010 |
| Reference male ejaculate<br>weight | 1 | 85.42 | 5.66 | 0.019 |
| Ejaculate dumping | 1 | 61.32 | 4.07 | 0.046 |
| Residual | 146 | 2202.11 |  |  |

A two-way linear model of variation in mating duration showed that males mated somewhat longer when mating as a female’s first (24.3 min) compared to second (21.2 min) mate (*F*_1,104_ = 4.24, *P* = 0.042), and that females with ablated hind-legs mated for longer (24.3 vs. 21.2 min)( *F*_1,104_ = 4.40, *P* = 0.038), although male genital treatment had no significant effect on mating duration (*F*_1,104_ = 0.02, *P* = 0.89). An analogous model of variation in male ejaculate weight showed that larger males transfer heavier ejaculates (*F*_1,103_ = 6.46, *P* = 0.013), while P1/P2, female treatment and male treatment had no significant effects (all *P* > 0.145). Interestingly, a model of female sperm dumping, simultaneously including both focal male body and ejaculate weight, showed that females dumped more ejaculate from relatively small males (*β′* = –0.019, SΕ_*β*_ = 0.004; *F*_1,102_ = 22.2, *P* < 0.001) with relatively large ejaculates (*β′* =1.29, SΕ_*β*_ = 0.09; *F*_1,102_ = 215.9, *P* < 0.001), but showed no effects of P1/P2, female treatment or male treatment (all *P* > 0.082).

A model of the number of scars in females, including the mating duration of both matings, revealed that the mating duration of the reference male (*F*_1, 100_ = 4.51, P = 0.036) was positively related to scarring and that females with ablated hind legs suffered fewer scars on average (136.2, SE = 7.7) than did females with intact hind legs (167.8, SE = 8.3) (*F*_1,100_ = 7.45, *P* = 0.007) but showed no effect of male treatment (*F*_1,100_ = 1.94, *P* = 0.166), suggesting that female resistance during copulation increases copulatory wounding. We failed to find any significant effects of any predictors on the area of scarring in females.

Female fecundity, i.e. the total number of eggs laid during our experiment, was positively associated with mating duration and tended to be positively related to ejaculate weight (Table 2). Notably, female fecundity was also affected by male genital treatment (Table 1), such that females laid fewer eggs if her focal male mate had ablated genital jaws (Fig. 5). This was true also in a reduced model, involving only AJ and Non males, where male treatment had a significant effect (*F*_1,96_ = 4.41, *P* = 0.038) while P1/P2, female treatment and their interaction had no significant effects (P > 0.074 in all cases).

**Table 2.**
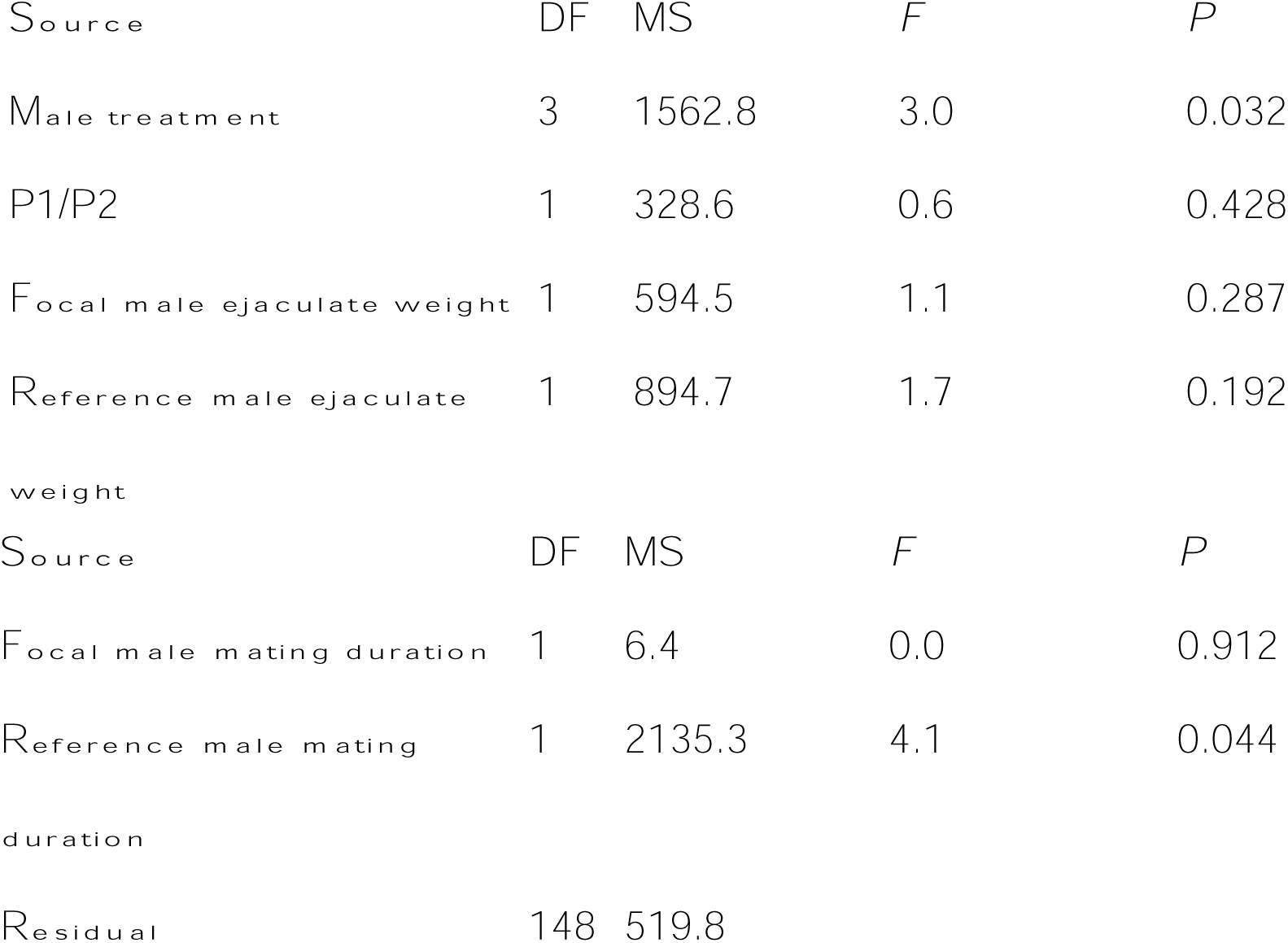
Analysis of variance of female fecundity.

| Source | DF | MS | F | P |
| --- | --- | --- | --- | --- |
| Male treatment | 3 | 1562.8 | 3.0 | 0.032 |
| P1/P2 | 1 | 328.6 | 0.6 | 0.428 |
| Focal male ejaculate weight | 1 | 594.5 | 1.1 | 0.287 |
| Reference male ejaculate weight | 1 | 894.7 | 1.7 | 0.192 |
| Focal male mating duration | 1 | 6.4 | 0.0 | 0.912 |
| Reference male mating duration | 1 | 2135.3 | 4.1 | 0.044 |
| Residual | 148 | 519.8 |  |  |

## Discussion

In contrast to the studies of the congener *C. maculatus* by Hotzy and Arnqvist (2009) and Hotzy et al. (2012), we found no significant effects of experimental ablation of genital spines on male fertilization success in *C. subinnotatus.* Instead, fertilization success was determined primarily by male ejaculate weight and the degree to which females dumped the ejaculate after mating. This suggests that females may affect male fertilization success, by differential uptake of male seminal fluid from relatively large males (Eberhard, 1996). Most importantly, we found that females laid fewer eggs following mating with males with ablated genital jaws, suggesting that this structure may ultimately function to stimulate female egg production more than female sperm use.

Hotzy et al. (2012) found that male seminal fluid is transported across the walls of the copulatory tract less rapidly in males with ablated genital spines and that such males suffer reduced fertilization success as a result. Our results suggest that the genital jaws of *C. subinnotatus* may also affect female uptake of male seminal fluid, although this may then be manifested as elevated female egg production in this species. We note that that male seminal fluid in seed beetles contains a very large number of proteins, some of which affect male fertilization success and others that affect female egg production (Goenaga et al., 2015; Yamane et al., 2015; Bayram et al., 2017). Needless to say, given everything else equal, male postmating reproductive success is elevated by an increase in female egg production (Arnqvist and Rowe, 2005).

Thus, our results provide support for a role of postmating sexual selection in the evolution of the genital jaws in *C. subinnotatus,* although the proximate mechanism is somewhat unclear and may differ somewhat from that seen in *C. maculatus* (Hotzy and Arnqvist, 2009; Hotzy et al, 2012). It is interesting to note that the fact that male ejaculate weight and the degree to which females dump ejaculate after mating determines male fertilization success is consistent with an important role for seminal fluid in mediating male postmating reproductive success also in *C. subinnotatus.*

We found that mating duration was positively associated with scaring in females, as has previously documented in *C. maculatus* (Crudgington and Siva-Jothy, 2000), and that females that were made unable to resist males by kicking suffered less scars. This shows that the physical act of resistance by females actually acts to aggravate the injuries they sustain during copulation. Ablation of genital spines decreases the amount of scarring suffered by female in *C. maculatus* (Hotzy and Arnqvist, 2009; Hotzy et al., 2012), but we found no significant effect of genital jaw ablation in *C. subinnotatus.* It is possible that our ablation treatment was too subtle to generate an effect on scarring in females strong enough for detection, in the face of rather extensive scarring in females caused by other genital spines.

Although it is certainly possible that the enigmatic genital jaws of male *C. subinnotatus* serves additional functions, we show here that spines on these jaws act to increase female egg production rate and are hence favored by postmating sexual selection. Whether sexually antagonistic coevolution has been involved in their elaboration is, however, less clear since sexual conflict relies on the demonstration of direct costs to females (Arnqvist and Rowe, 2005; Fricke et al., 2009). An elevation of egg production rate may come at a nest cost to females (Arnqvist and Rowe 2005), but this depends on how increased the rate of egg production trades-off with other fitness components (Rönn et al., 2006). Similarly, our experiments suggest that the increased scarring caused by the genital jaws is relatively marginal and direct costs to females may thus be minor. Additional studies are required to further clarify the role of the genital jaws in *C. subinnotatus* and to assess the degree to which this remarkable structure is detrimental to females.

